# Seizures induce c-fos expression in a subset of astrocytes, termed fostrocytes, that dampen subsequent seizures

**DOI:** 10.64898/2026.09.01.748637

**Authors:** Madison J. Failor, Bradley P. Bork, Jennifer M. San Pietro, Aleksandra Maciejczuk, Kendall M. Elmer, Corinne Lile-King, Mariia Oliinyk, Junguhk L. Choi, Andrew H. Huang, Dominik J. Tabor, Ronald P. Gaykema, Edward Perez-Reyes

**Affiliations:** Biomedical Sciences Graduate Program, University of Virginia, Charlottesville, VA 22903, USA; Department of Pharmacology, University of Virginia, Charlottesville, VA 22903, USA; Brain Institute, University of Virginia, Charlottesville, VA 22903, USA

**Keywords:** astrocytes, c-fos, spontaneous seizures, temporal lobe epilepsy

## Abstract

**Objective:** The original goal was to map neuronal circuits activated by spontaneous seizures in models of temporal lobe epilepsy. Studies used the c-fos driven TRAP2 system, which has been used successfully to label neurons after seizures. Unexpectedly, astrocytes were also labeled, then shown to express c-fos in a sustained manner after seizures. The role of these so-called fostrocytes in spontaneous seizures was studied using novel Cre-dependent AAVs.

**Methods:** Studies used a homozygous mouse line produced by crossing TRAP2 Cre-driver mice with the Ai9 Cre-reporter line. Seizures were induced using electrical stimulation, kainic acid, or pilocarpine. Cre-dependent AAVs used the GFAP promoter to drive expression of either the catalytic A chain of diphtheria toxin (DTA), GFP, or empty vector. Spontaneous seizures were continuously recorded using EEG.

**Results:** Discrete seizures in naïve mice evoked transient c-fos expression in all astrocytes, while status epilepticus evoked a higher and sustained level of c-fos expression. Timing the activation of the TRAP2 system allowed specific labeling of the subset of astrocytes with high c-fos expression. These cells were colocalized with established astrocyte markers, showed reactive astrocyte morphology, and were selectively labeled by GFAP-driven Cre-dependent AAVs. Ablation of fostrocytes in spontaneously seizing mice increased seizure frequency.

**Significance:** Next-generation sequencing studies have revealed a great diversity of astrocyte subtypes, identifying clusters with up-regulated c-fos expression in patients with neurological disorders. In conclusion, these studies suggest that therapies that augment the activity of c-fos expressing astrocytes would have anti-seizure activity.

**Key points:**

➢ Discrete seizures triggered a transient expression of c-fos in all astrocytes, while status epilepticus triggered its prolonged expression in a subset of astrocytes.
➢ The TRAP2 system expresses inducible Cre recombinase controlled by the Fos gene, providing genetic access to astrocytes that express c-fos at high levels.
➢ Cre-dependent delivery of a diphtheria toxin selectively ablated these fostrocytes and exacerbated spontaneous seizures.
➢ Novel therapies that augment the activity of fostrocytes are predicted to reduce seizure burden.

## 1. INTRODUCTION

Epilepsy is a common neurological disorder that affects over 50 million people worldwide. Despite significant improvements in efficacy and lowered side effects, seizures remain poorly controlled in about one-third of epilepsy patients. One of the most common treatment-resistant epilepsy syndromes is temporal lobe epilepsy (TLE). The seizure onset zones in TLE reside in hippocampal and parahippocampal brain regions. The hippocampus is particularly vulnerable to seizures, and human TLE can be caused by a wide variety of insults such as status epilepticus (SE), febrile seizures, dysplasias, infection, stroke, and traumatic brain injury.^1^

Current animal models of TLE begin by inducing SE, which can be triggered electrically by placing a stimulating electrode in the hippocampus or with chemoconvulsants that activate hippocampal circuitry, such as pilocarpine and kainic acid.^2^ Evidence that these animal models are externally valid proxies for human TLE include EEG patterns, tonic-clonic motor seizures, hippocampal sclerosis, and localization of the seizure focus to the hippocampal formation.^3^ Our approach is to develop gene therapies that target seizure circuits activated by spontaneous limbic seizures using TRAP2 mice crossed with Ai9 reporter mice.^4, 5^ The *Fos* gene in these mice was engineered to express a Cre recombinase that can be activated by tamoxifen and/or 4-hydroxy-tamoxifen (4OHT). TRAPing of an active cell requires the simultaneous expression of Cre recombinase and exposure to 4OHT. The *Fos* gene is tightly regulated, being turned on and off in minutes, and this rapid turnover extends to the c-fos mRNA and protein.^6, 7^ For example, Peng and Houser showed that hippocampal c-fos protein expression increases rapidly after a seizure, peaking between 30 and 60 minutes, and then decaying rapidly by 120 minutes.^8^ In contrast, Cre recombinase is a relatively stable protein in neurons.^9^ Therefore, we sought to determine the longest duration after an evoked seizure that 4OHT effectively traps activated cells. These experiments led to the discovery that seizures activate a subset of glial cells. This study established that these cells are astrocytes and determined the time course of their activation after seizures. We developed novel Cre-dependent and astrocyte-specific AAV tools to ablate astrocytes based on their expression of the *Fos* gene. We then used these AAVs to interrogate the role of these c-fos-expressing astrocytes in various mouse models of TLE, including electrically-kindled and chemoconvulsant models.

## 2. METHODS

### 2.1 Animals

Experimental procedures involving mice conformed to institutional policies and ARRIVE guidelines. ^10^ Mice had ad libitum access to food and water and were kept in vivarium facilities with controlled temperature, humidity, and 12 h light/dark cycle. Both male and female mice were used in each experiment at equal ratios. We used TRAP2^4^ mice (Fos^2A-iCreERT2^, #030323, Jackson Laboratories) that were crossed with Ai9^5^ tdTomato reporter mice (B6.Cg-Gt(ROSA)26Sor^tm9(CAG–tdTomato)Hze^/J, #007909, Jackson Laboratories). Mice were crossed for over 10 generations and are on a C57BL/6 background. Mice that were homozygous for both alleles were crossed with CD1 mice for the pilocarpine studies (#034608 Jackson Laboratories).

### 2.2 Surgeries

Detailed methods describing the surgeries to implant the EEG recording headsets have been described previously.^11, 12^ AAVs were bilaterally injected along a vertical axis extending from -4.6 to -2.4 mm below bregma targeting the center of the hippocampus (3.6 mm lateral and -3.3 posterior from bregma). Pain was minimized with bupivacaine and ketoprofen. We used the following AAVs packaged in serotype 5 particles (Vectorbuilder): 1) GLFFlxGSDTA (GFAP driven split diphtheria toxin A, GSDTA); 2) GFDiGFP-H2 (GFAP-driven GFP tagged with H2B); and 3) GFAPNCtrl (GFAP promoter empty payload). Supplemental Fig. 1 details the AAV constructs and the injection scheme. All research personnel were blinded to the AAV injected.

### 2.3 Seizure Induction and EEG Recording

Table 1 lists the protocols used to induce seizures in TRAP2 mice and how seizures were monitored. Identification studies used naive mice and seizures were scored by multiple observers in real time. Studies on spontaneous seizures in chronically epileptic mice used two methods to induce epilepsy: hybrid kindling and pilocarpine. Hybrid kindling combines escalating systemic doses of kainic acid^13^ (KA, 10-5 mg/kg) followed by electrical stimulation until fully kindled.^14^ We used an improved version of the pilocarpine model where mice are treated while on the EEG rig.^12^ The protocol includes pretreatment with N-methyl-scopolamine (1.75 mg/kg) 20 min before pilocarpine (159 mg/kg), and then after 1 h of SE, treatment with diazepam (10 mg/kg). Mice were then fed soft rodent chow supplemented with Nutrical for at least 3 days. To induce iCreERT2 recombination, we used 4OHT (40 mg/kg, Med-Chem Express), an analog of tamoxifen that activates iCreERT2 at 100-fold lower concentrations than tamoxifen.^15^ Notably, 4OHT was dissolved in DMSO, diluted 2-fold with Cremophor EL (MilliporeSigma), and then diluted 5-fold into saline.^12^ EEG/video monitoring (continuous 24 hours/day, 7 days/week) was performed from the induction of evoked seizures to the end of the experiment.^12^ EEG recordings were used to estimate the duration of SE and subsequent seizure frequency using published criteria.^16^ The EEG was read by two researchers who were blind to the experimental procedure.

**Table 1.** seizure models used in this study.

|  | No EEG, discrete GTCS | EEG, SRS | Location |
| --- | --- | --- | --- |
| Kainic acid | + |  | Fig. 1 |
| Pilocarpine | + |  | Fig. 1 |
| Hybrid kindled |  | + | Fig. 3 |
| Pilocarpine |  | + | Fig. 2, 5, & 6 |

Abbreviations: GTCS, generalized tonic-clonic motor seizure; SRS, spontaneous recurring seizures, Hybrid kindled, combination of kainic acid and electrical kindling.

### 2.4 Immunohistochemistry

Methods used for antibody staining have been described previously.^17^ We used the following primary antibodies to stain 40 µm horizontal brain slices: mouse anti-S100b (RRID: AB_477499), goat anti-sox9 (RRID: AB_2194160), chicken anti-GFAP (RRID: AB_2313547), rabbit anti-Fos (RRID: AB_2106755), and rabbit anti-dsRed (RRID: AB_10013483). The corresponding secondaries used were goat anti-mouse Alexa Fluor 647 (RRID: AB_2338906), donkey anti-goat Alexa Fluor 647 (RRID: AB_2340436), goat anti-chicken Alexa Fluor 647 (AB_2337392), donkey anti-rabbit Alexa Fluor 488 (RRID: AB_231358), and goat anti-rabbit Alexa Fluor 568 (RRID: AB_143157).

### 2.5 Image Acquisition and Analysis

We used stereological techniques as described previously.^17^ Imaging was performed on an Olympus IX81 confocal microscope equipped with a Hamamatsu Orca 4.0 camera. Acquisition and analysis were performed with cellSens software. Cell count analysis was performed on images acquired with a 20x objective. Morphology analysis was performed on Z-stack images acquired with a 60x objective. Neurolucida 360 was used to analyze 60x images, trace the processes of astrocytes, and perform Sholl and convex hull analysis.

### 2.6 Statistics

Findings were compiled in Excel spreadsheets and then imported into GraphPad Prism (v. 11) for statistical analysis and figure creation. All data sets were checked for normality, and an appropriate ad-hoc test was chosen. The statistical tests used are defined in the figure legends. A P value of 0.05 was used to determine statistical significance. In graphs with multiple comparisons, we only show the P value if it was below 0.05.

## 3. RESULTS

### 3.1 Discovery and identification of seizure-activated astrocytes in TRAP2 mice

Our overall goal is to develop anti-seizure gene therapies that target epileptic circuits.^18^ We study these circuits using the TRAP2 system.^4, 19, 20^ Our studies began with a time course experiment to determine how long after a seizure we could still “TRAP” c-fos-expressing cells with 4OHT. Seizures were induced in TRAP2 mice using kainic acid and 4OHT was injected at various time points thereafter (Fig. 1A). In most mice, KA caused 3-5 discrete seizures accompanied by convulsive motor seizures. We waited 2 weeks after 4OHT administration to allow for tdTomato expression and then fixed the brain and prepared horizontal slices. Images of the hippocampus from mice TRAPed within 1.5 h of the seizure showed strong tdTomato expression in neurons (Fig. 1B). In contrast, images of mice TRAPed after 3 h showed lower levels of neuronal activation. The decay of the neuronal signal that we observed followed a similar time course to that previously reported by DeNardo and colleagues^4^ (Fig. 1C). Surprisingly, images from mice TRAPed 24 h after KA displayed tdTomato positive (tdT+) cells with a glial morphology. Interestingly, these tdT+ cells activated at much later time points than neurons, peaking 24 h after a seizure (Fig. 1C). To identify these tdT+ cells, we performed immunohistochemistry with antibodies for the established astrocytic markers GFAP, S100B, and Sox9.^21^ The results of these colocalization studies showed that the majority of tdT+ glia were stained with all 3 antibodies, identifying them as bona fide astrocytes (Fig. 1D, E). We dubbed these TRAPed tdT+ astrocytes, fostrocytes.

**Fig. 1.**
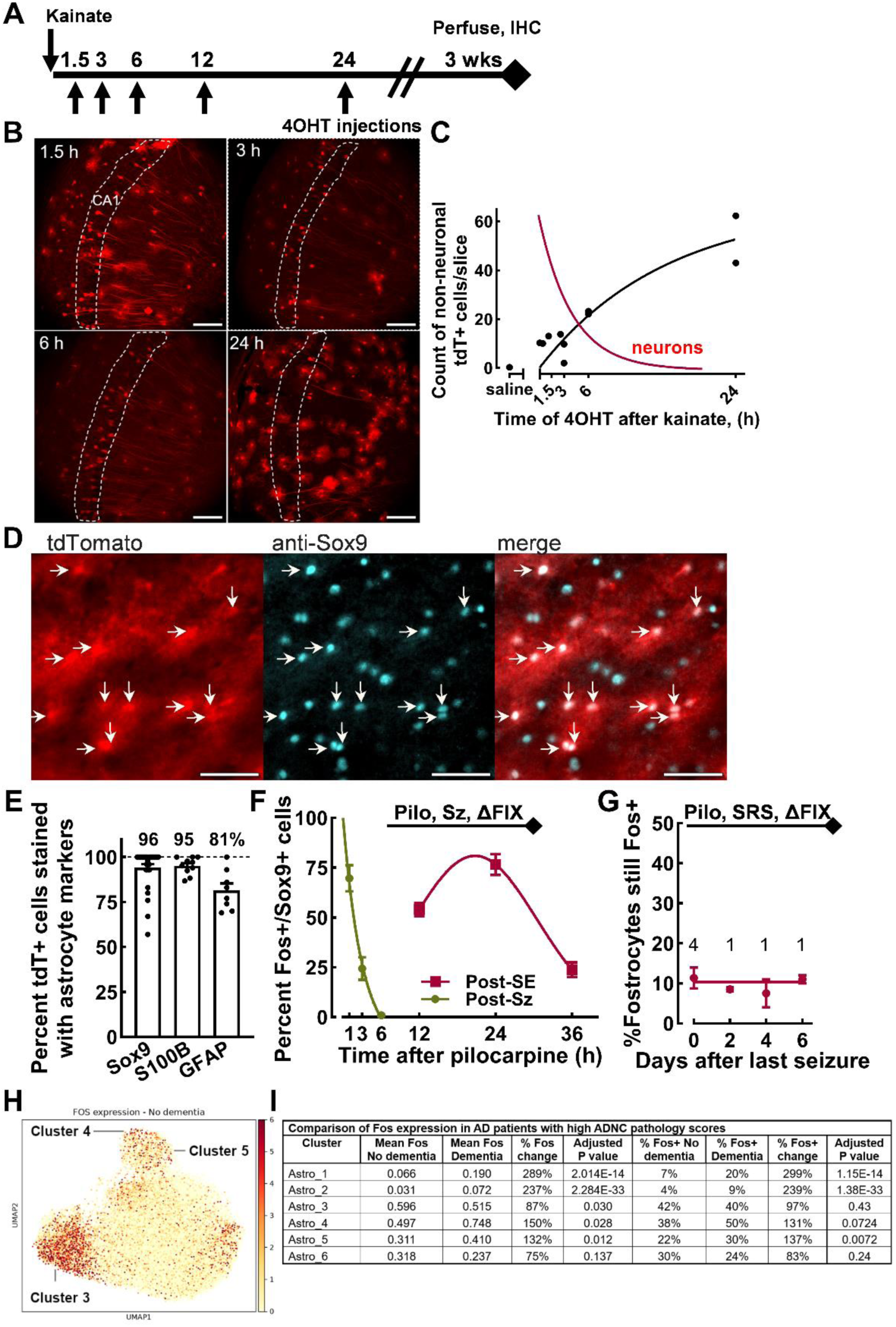
Discovery and identification of seizure-activated astrocytes in TRAP2 mice. **A,** Schematic of the timeline for TRAPing cells activated by seizures with single 4OHT injections. **B,** Representative images of horizontal brain slices illustrating cells TRAPed in CA1 either 1.5, 3, 6, or 24 h after kainic acid (20x images; scale bar, 100 μm). Dotted lines outline the CA1 pyramidal cell layer. **C,** Plot of the time course of TRAPing cells after kainic acid induced seizures (each point represents the average tdT+ count per mouse, N=11). Curves represent single exponential fits to the data. Red line represents a single exponential fit to the time course of TRAPing neurons. **D-E,** Colocalization of the tdT+ non-neuronal cells with validated astrocyte markers. **D,** Representative multi-channel images of tdTomato and anti-Sox9 signal. The secondary antibody for Sox9 staining was Alexa-647 and was pseudo-colored to create a white signal in the merged panel (white arrows; 20x images; scale bar, 10 μm). **E,** Quantification of the colocalization of tdT+ cells with Sox9, S100B, and GFAP (Sox9, N=30 images, 208 cells, from 2 mice; S100B, N=10 images, 911 cells, from 5 mice; GFAP, N=4 images, 208 cells, from 1 mouse). **F-G,** Time course from two separate experiments where seizures were induced by pilocarpine. **F,** Time course of anti-fos immunoreactivity in astrocytes stained by anti-Sox9. Pilocarpine either triggered discrete seizures (green) or SE (red). The group with discrete seizures was used for 1, 3, and 6 h points (N=3, 3, 2 mice, respectively), while the post-SE group was used for 12, 24, and 36 h points (N=2, 3, 2 mice, respectively). Early time points were fit to a single exponential (r^2^=0.84), while later points were fit to a Gaussian (r^2^=0.86). **G,** Fostrocytes that remain Fos+ after 5 weeks of SRS. Mice were TRAPed while spontaneously seizing, and then later fixed and stained with anti-Fos antibody. Post-hoc analysis of the seizure data allowed us to calculate the days after their last seizure that the brains were fixed. Four of the mice seized on the last day of the experiment, while there was one mouse for the other time points. Results include counts from each hemisphere. Data were fit to a horizontal line. **H,** Heat map of Fos RNA expression in patients with No Dementia. Data from the single-cell profiling dataset of the Seattle Alzheimer’s Disease Brain Cell Atlas (SEA-AD^43^). Results from our analysis of the single-nucleus transcriptomic data of the human middle temporal gyrus, which was generated using 10x 3′ v3 and 10x Multiome assays. Data was accessed July 2026 through the CELLxGENE Discover SEA-AD Collection and the Registry of Open Data on AWS. This dataset contains 70,009 annotated astrocyte nuclei from donors classified as normal or having dementia. **I,** Fos expression and Fos-positive astrocytes in high-ADNC cases with and without dementia. Mean Fos expression is the mean log1p-transformed expression normalized to 10,000 counts per nucleus. Fos-positive nuclei were defined as normalized Fos expression >0.1. Fos+ percentages were compared using two-sided two-proportion z-tests, with p-values adjusted using the Benjamini-Hochberg method across the six astrocyte-cluster comparisons. Analysis was performed using Scanpy v1.11.5.

Fos is an immediate early gene transcription factor activated by cAMP and Ca^2+^ second messenger signaling, and hence, is an accepted proxy for neuronal activity. However, the roles of c-fos in astrocyte physiology are only a recently emerging theme.^22, 23^ Our findings raised many questions. Do all astrocytes express c-fos after a seizure? Are all c-fos+ astrocytes capable of being TRAPed by 4OHT? To address these questions, we turned to the pilocarpine model of TLE using our improved protocols to induce both discrete seizures and SE.^12^ The results shown in Fig. 1F were obtained from mice that were treated with pilocarpine and then fixed with formalin soon after. This experiment does not include 4OHT. Immunohistochemistry was performed using antibodies to c-fos and Sox9 with colocalization as determined as above. Interestingly, discrete seizures triggered a transient expression of c-fos in virtually all astrocytes, while prolonged SE triggered a second wave of c-fos expression in a subset of astrocytes (Fig. 1F). We then asked: how long after a spontaneous seizure is c-fos expressed in a TRAPed fostrocyte? To address this question, we studied c-fos expression at the end of a chronic epilepsy study that will be presented later. In this experiment we recorded spontaneous seizures in TRAPed mice, and post-hoc identified mice that did not have a seizure in the days before formalin fixation. These studies show that ∼10% of TRAPed fostrocytes continue to express c-fos for many days after seizures (Fig. 1G). Taken together, fostrocytes can be operationally defined as a subset of astrocytes that respond to seizures with a state change characterized by a slow and prolonged activation of *Fos* gene expression, which reaches a level high enough to induce Cre recombination of floxed tdTomato in TRAP2 mice.

Since seizures in people with dementia are associated with AD pathology, such as hippocampal atrophy,^24^ we wondered if these patients had altered *FOS* gene expression. Data mining of the Seattle Alzheimer’s Disease Brain Cell Atlas (SEA-AD) revealed an upregulation of *FOS* and *JUN* in multiple astrocyte clusters. Fig. 1H shows a heat map of Fos expression in the superclusters identified in control subjects. Fig. 1I compares Fos RNA expression in all patients with high Alzheimer’s Disease Neuropathologic Change (ADNC) scores,^25^ in terms of mean expression and percent of cells in each cluster that are Fos+.

### 3.2 Fostrocytes are activated after spontaneous recurring seizures in 2 models of TLE

To establish that fostrocytes are activated by spontaneous seizures regardless of the animal model, we compared fostrocyte counts in mice that were either hybrid-kindled or pilocarpine treated (Fig. 2). A key difference in these models is that SE is absolutely required to induce spontaneous seizures in the pilocarpine model. In both experiments there were mice that did not develop spontaneous recurring seizures (No SRS). Importantly, 4OHT was injected during the chronic phase when most mice have SRS. Fostrocyte counts in mice with “No SRS” were low to non-detectable in both hybrid kindled and pilocarpine-treated mice (Fig. 2B & E). In contrast, fostrocyte counts were considerably higher when 4OHT was injected into mice with SRS. In both models, fostrocytes were observed primarily in the hippocampus, and their abundance followed the rank order CA1 ≥ CA3 > dentate gyrus. Notably, pilocarpine-treated mice with SRS had over two times more fostrocytes in the hippocampus than hybrid-kindled mice with SRS. Lastly, naïve mice with no evoked or spontaneous seizures had no fostrocytes whatsoever (data not shown). These experiments confirm that seizures are required to induce the TRAPing of fostrocytes, and that this occurs after spontaneous seizures in epileptic mice.

**Fig 2.**
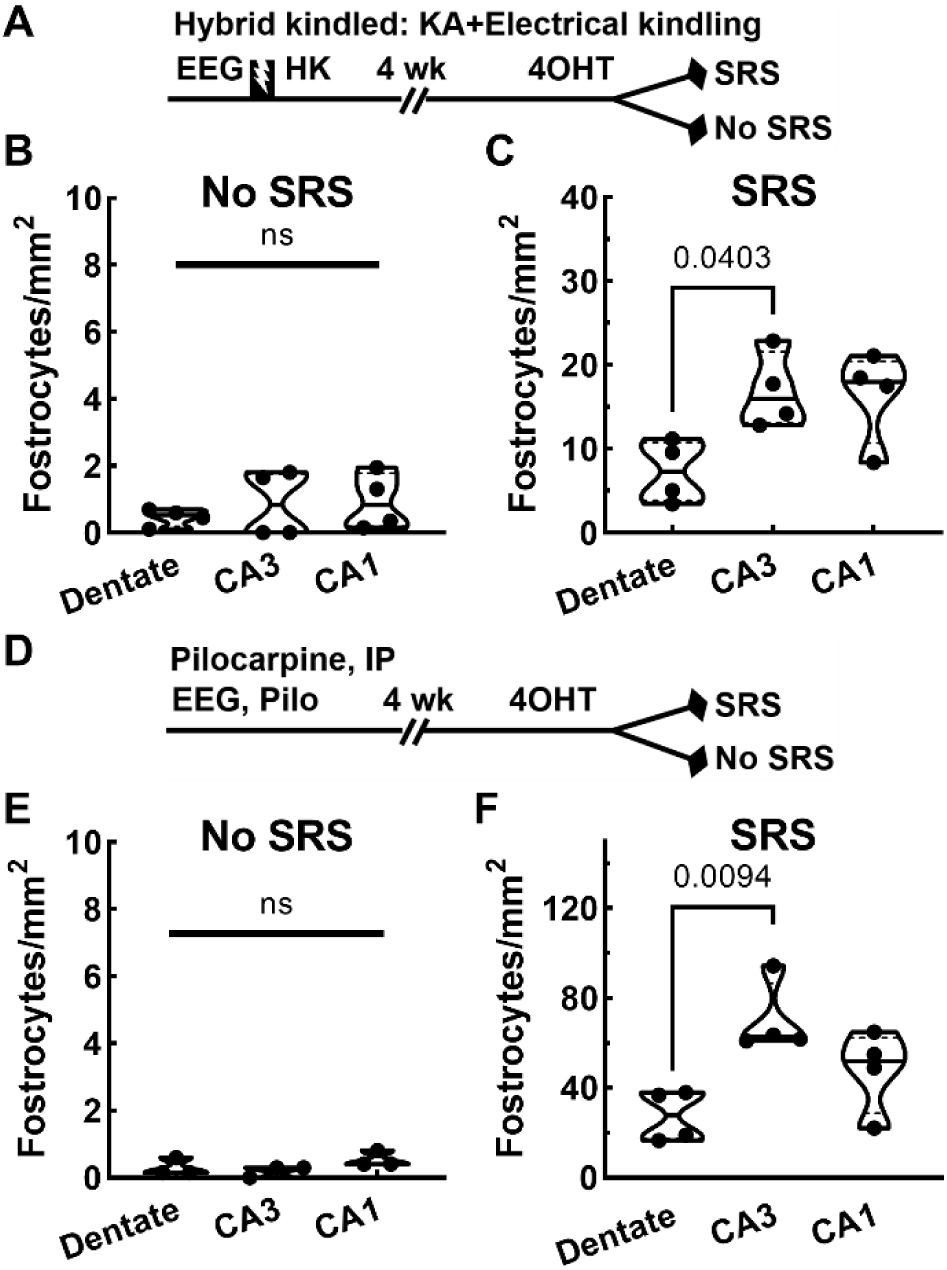
Fostrocytes are activated after spontaneous seizures in 2 models of TLE. **A,** Schematic of experimental timeline for mice in B and C, where mice were treated with kainic acid and then electrically kindled until criterion for fully kindled (>5 GTCS; Hybrid-Kindled, HK). In this protocol, only half the mice develop spontaneous recurring seizures (SRS). 4-Hydroxy-tamoxifen was injected after 4 weeks of EEG recording. **B,** Fostrocyte counts in slices from hybrid-kindled mice with no spontaneous seizures (n=4 mice, each point is the median count per mouse from 4 depths). **C,** In contrast, fostrocytes were observed all over the hippocampus in mice that developed spontaneous seizures. Fostrocytes were particularly concentrated in CA3 and CA1 fields over the dentate gyrus. **D,** Schematic of experimental timeline for inducing spontaneous seizures in mice using pilocarpine. In this experiment 3 mice did not develop SRS, while 4 mice developed SRS. **E**, Fostrocyte counts in slices from pilocarpine mice with no SRS were very low in all subfields. As observed with the HK model, fostrocytes were abundant in the hippocampus, with higher density again found in CA fields. Statistical analysis by one-way ANOVA, and only statistically significant comparisons are shown.

### 3.3 Fostrocytes adopt a reactive astrocyte morphology

Astrocytes are known to change morphology in many neurological disorders. Therefore, we examined the morphology of fostrocytes in two post-SE models. Figure 3 compares the morphology of astrocytes in naïve TRAP2 mice to tdT+ fostrocytes and tdT-astrocytes from pilocarpine and kainic acid treated epileptic mice. Morphology differences were strikingly apparent when comparing tracings of fostrocytes to astrocytes in naïve mice. Sholl analysis was performed to assess branch complexity differences, and it was found that fostrocytes have significantly fewer branches than astrocytes from naïve samples (Fig. 3B). The average branch diameter was significantly higher in fostrocytes (Fig. 3C), while branch volume was significantly lower compared to naïve astrocytes (Fig. 3D). In contrast, the morphology of tdT+ fostrocytes was virtually identical to adjacent tdT-astrocytes in the same slice (Fig. 3D-F). These results were also replicated in slices from hybrid-kindled mice, with fostrocytes adopting a reactive phenotype (Fig. 3B-D).

**Fig. 3.**
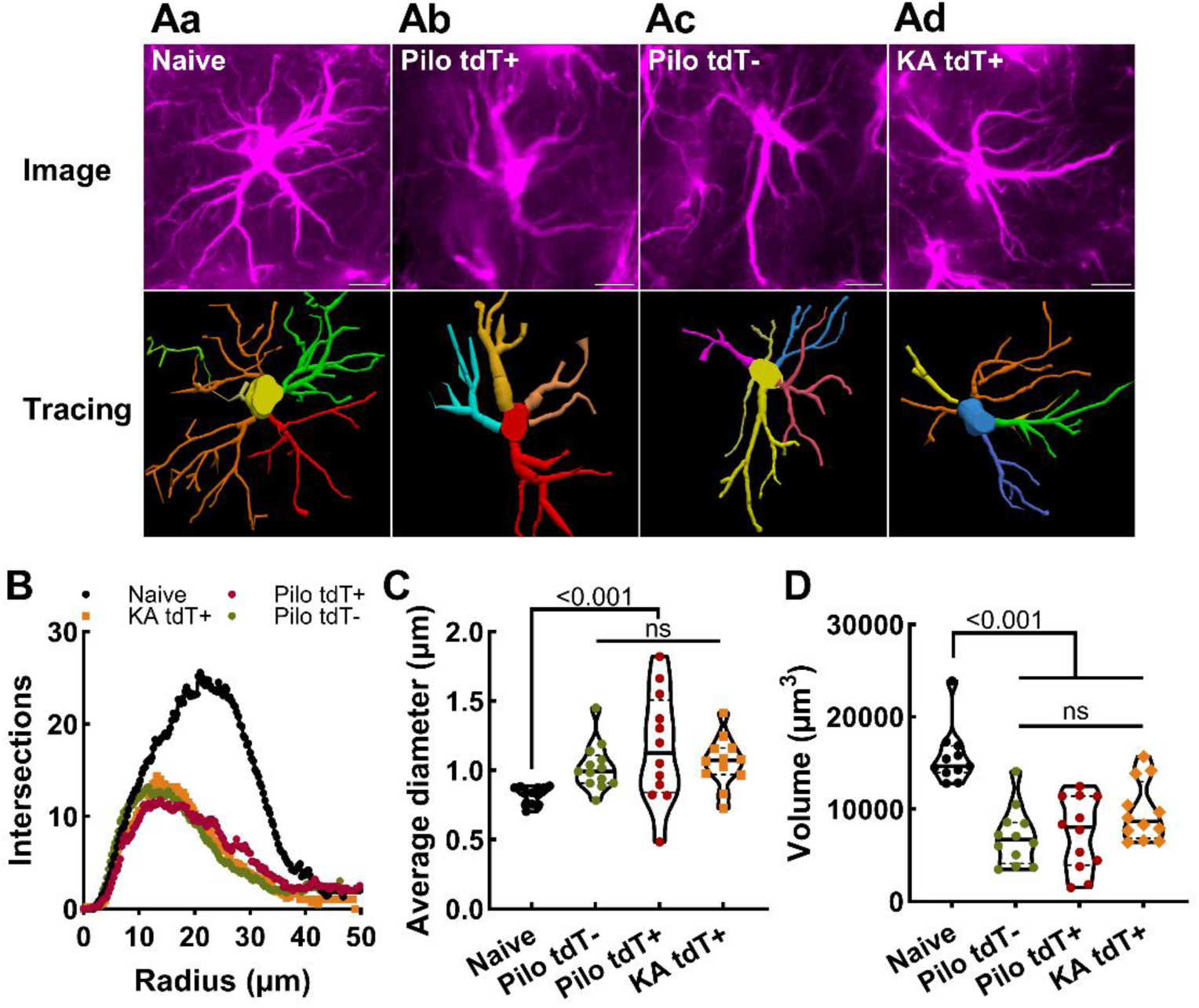
Fostrocytes adopt a reactive astrocyte morphology. **A,** Representative images of fostrocytes and after Neurolucida tracing. To visualize astrocyte processes, brain slices were stained with both anti-GFAP and anti-S100B using a common secondary antibody and then imaged in Z-stacks using a 60X objective. Results are shown for: **Aa**, an astrocyte from a naïve TRAP2 brain; **Ab**, a tdT+ fostrocyte from an epileptic pilocarpine-treated brain; and **Ac**, a tdT-astrocyte in an epileptic pilocarpine-treated brain. **B,** Sholl analysis of these 3 groups, and including results from tdT+ fostrocytes in the hybrid kindled model (n=12 cells, each point represents the average intersection at a given radius). The Neurolucida tracings were used to determine the average diameter of astrocyte processes (**C**) and their volume (**D**). In both cases, naïve astrocytes showed thinner processes that extended over a larger volume than astrocytes found in brains from epileptic mice, while astrocytes from mice with spontaneous seizures had fatter processes and occupied a smaller volume. Statistical analysis by one-way ANOVA. Each point represents a cell (n=12/group) imaged from 2-3 mice per group.

### 3.4 Validation of novel Cre-dependent AAVs

Based on the finding that reactive astrocytes can be neurotoxic,^26^ we hypothesized that fostrocytes played a causative role in epilepsy. To test this hypothesis, we engineered novel Cre-inducible AAVs to deliver the catalytic A chain of diphtheria toxin (DTA). This chain catalyzes the ADP-ribosylation of ribosomal elongation factor-2, thereby stopping protein synthesis and inducing apoptotic death.^27^ Cre-dependence relies on the use of interwoven Cre binding sites called DIO or FLEX cassettes (Fig. 4A).^28^ The FLEX cassette contained a split DTA (GSDTA), where the DTA coding sequence was split into two fragments with one fragment being inverted in the FLEX cassette (Fig. 4B, Supplemental Fig. 1). The FLEX cassette used lox66 and lox71 Cre binding sites to lower non-Cre mediated expression.^29^ We used the GFAP promoter fragment, gfaABC1, to target delivery to astrocytes.^30^ Our AAV constructs also included mir-124 binding sites, which were demonstrated to improve targeting to astrocytes.^31^ In these studies, over 90% of the cells expressing GFAP-DIO-GFP-H2 were Sox9+ astrocytes (Fig. 4B). The ability of GSDTA to ablate GFP-H2-labeled astrocytes was tested in epileptic mice treated with 4OHT after a seizure. These results show GSDTA was highly effective, ablating over 90% of GFP-H2+ cells (Fig. 4C). Leak expression from GSDTA was assayed using naive C57BL/6 mice that do not express Cre recombinase (Fig. 4D). We used unilateral injection of GSDTA and then compared Sox9+ astrocyte counts between the injected and non-injected hemispheres. We found no statistically significant differences between the two sides.

**Fig. 4.**
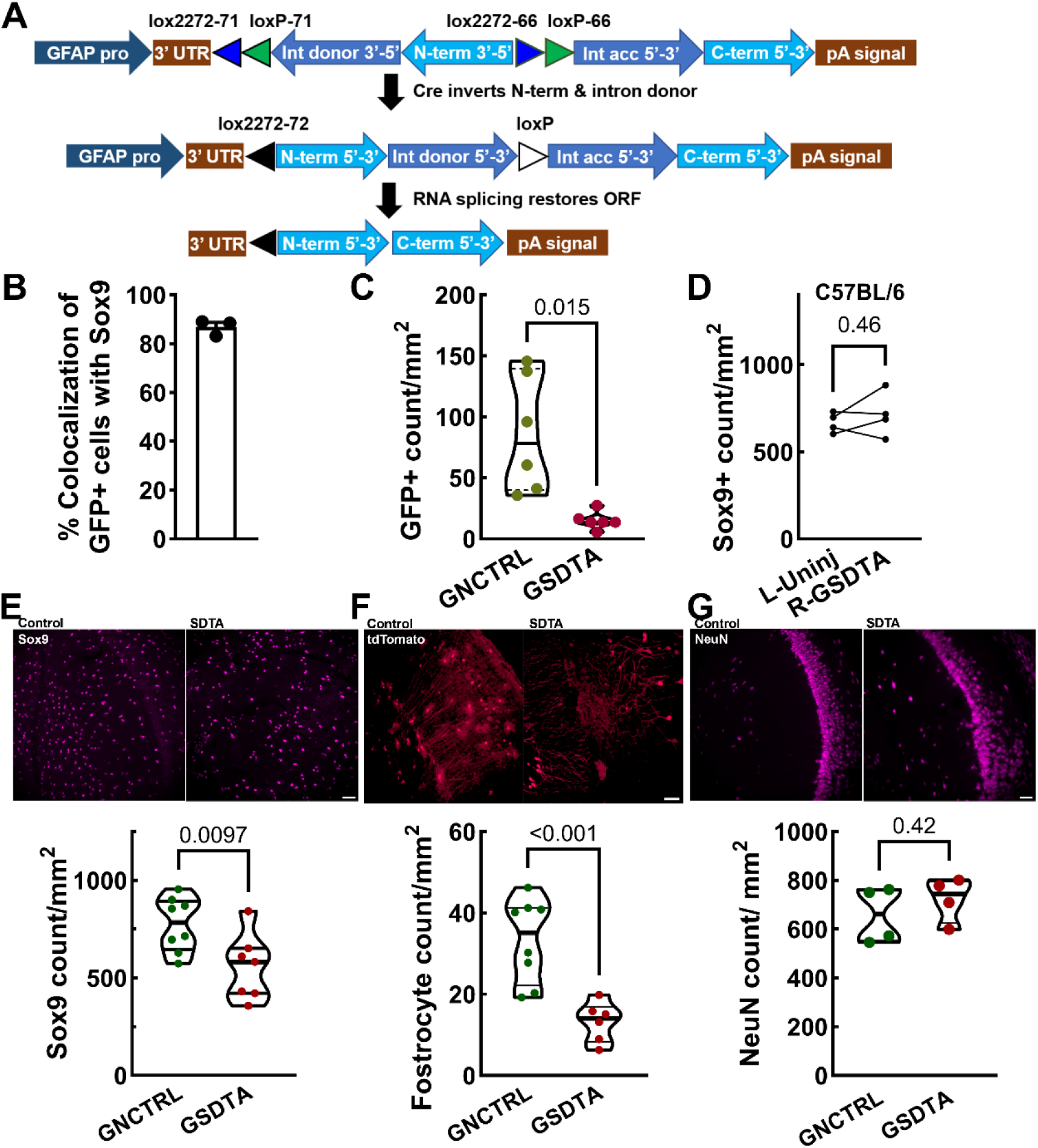
GFAP-FLEX2-GSDTA specifically ablates astrocytes without off-target ablation of neurons in TRAP2 mice. **A,** Schematic of the AAV construct before and after Cre recombination. Key details of the construct include: 1) use of the 2.2-kb GFAP promoter fragment; 2) mutant Cre binding sites (lox66 and lox71, FLEX2); 3) attenuated split Diphtheria Toxin A chain (GSDTA, G128D) where the N-terminal fragment was inverted and flanked by the FLEX2 excision cassette; 4) mIR-142 binding sites to inhibit expression in neurons; and 5) split introns to remove residual lox sites from the coding region after RNA splicing. **B,** Selectivity of the GFAP promoter was verified by colocalizing the green fluorescent signal from GFAP.DIO.GFP-H2 and anti-Sox9 antibody staining (each point is the average determination from 3 slices for 3 naïve TRAP2 mice. **C**, Off-target expression in the absence of Cre recombinase was assayed after unilateral injection of GSDTA into C57/BL/6 mice, followed by anti-Sox9 staining and counting astrocyte nuclei. The injection of GSDTA was inferred by expression of GFP from AAV-GFAP.DIO-GFP-H2. Statistical analysis by paired t-test, where each pair of points represents the results from 1 brain slice (N=9 slices, N=3 mice). **D**, Target engagement studies showing near complete ablation of GFP-tagged astrocytes using GSDTA. Statistical test Welch’s t-test (N=6 mice per group). **E**, Representative images and analysis of anti-Sox9 antibody-stained astrocytes in epileptic GNCTRL and GSDTA groups (20x images, scale bar, 100 µm). **F,** Representative images and analysis of anti-tdTomato antibody stained fostrocytes in epileptic mice injected with either GNCTRL or GSDTA mice (20x images, scale bar, 100 µm). Images and results are from the CA1 region. The results shown in panels G and H are the average per animal from 2 depths using an N=4 mice per group. Statistical analysis by Welch’s t-test. **G,** Representative images and analysis of anti-NeuN antibody staining of neurons in epileptic GNCTRL and GSDTA groups. Comparison of the NeuN stained CA1 neuron counts using 4 mice per group. Statistical analysis by unpaired t-test.

We next measured the effects of GSDTA on total astrocyte count, fostrocyte count, and neuronal count in epileptic mice (Fig. 4E-G). Astrocyte marker Sox9 was used to count astrocytes in the CA1 of the hippocampus of the GSDTA and GNCTRL groups (Fig. 4E). We used the tdTomato fluorescent signal to specifically compare the count of fostrocytes in GSDTA and GNCTRL mice in the CA1 region. Consistent with the results shown in Fig. 4E, the GSDTA group had significantly fewer astrocytes (Fig. 4E) and fostrocytes (Fig. 4F). Importantly, GSDTA did not significantly ablate neurons (Fig. 4G). These experiments validated the use of GSDTA to selectively ablate fostrocytes in epileptic TRAP2 mice, with no measurable effect on neurons or in non-Cre C57BL/6 mice.

### 3.5 Ablation of fostrocytes with GSDTA increases seizure frequency in TRAP2 mice

The experimental design included a single surgery to inject AAV and implant the EEG recording headset, 3 weeks of recovery, and then pilocarpine treatment while measuring the EEG response (Fig. 5A). Activation of Cre recombination by 4OHT was performed 24 hours after SE. We compared the response between two groups: the control group injected with GFAP-DIO-GFP-H2 plus GNCTRL AAVs, and the experimental group injected with GFAP-DIO-GFP-H2 plus GFAP-FLEX2-GSDTA (Supplemental Fig. 1D). A key advantage to this injection scheme is that toxic effects of fluorescent proteins are controlled,^32^ rather than compared to the treatment arm. A second advantage is that total AAV load can be normalized between injections using the GNCTRL AAV. Our studies used an EEG rig capable of recording 16 mice 24 h/day, 7 d/wk, allowing for 8 mice per cohort. Pilocarpine was typically dosed at 159 mg/kg and triggered SE in most mice. Importantly, the duration of SE was not different between GNCTRL and GSDTA groups (Fig. 5B).

**Fig. 5.**
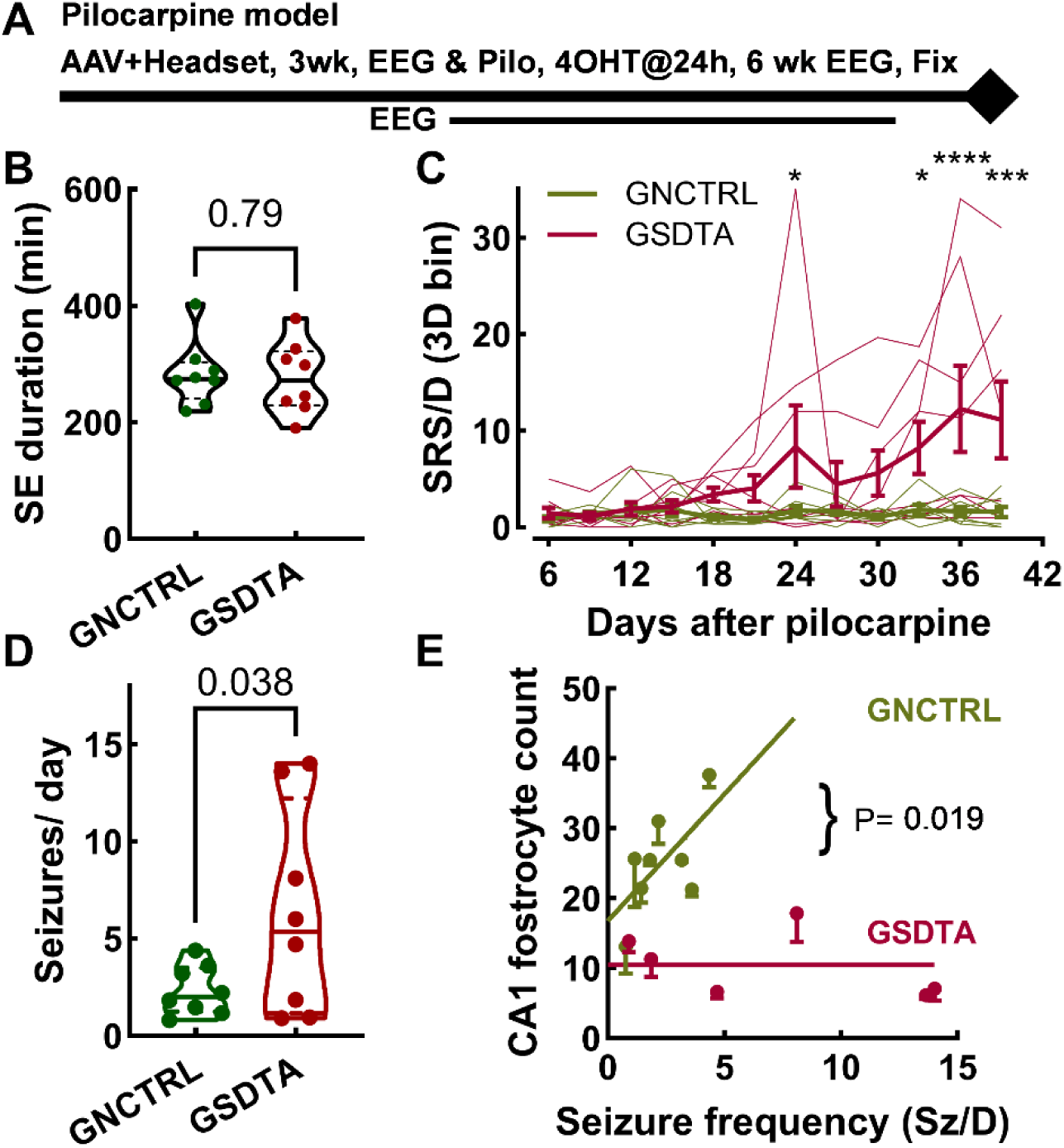
Ablation of fostrocytes increases seizure frequency. **A,** Schematic of the experimental timeline, which includes surgery for both AAV injection and EEG headset implantation. **B,** Duration of pilocarpine-induced SE was similar between GNCTRL and GSDTA groups. Statistical analysis by unpaired t-test (n=8 per group, each point represents results from one mouse). Treatment was performed while monitoring EEG, which allowed accurate measurement of when mice entered SE and when to inject diazepam. **C,** Plot of the seizure frequency versus time from pilocarpine treatment. Due to the high day-to-day variability in seizure frequency, the data were binned in 3 day epochs. Thin lines represent the seizures measured in each mouse, while thick lines show the average +/- sem of each group. Mice injected with GSDTA had significantly more seizures/day, particularly after 3 weeks of recording. Statistical analysis by 2-way ANOVA and Fisher’s Least Significant Difference test. Asterisks represent the following P values: *, P<0.05; ***, P<0.001; and ****, P<0.0001. **D,** Seizure frequency over the entire EEG recording period. The GSDTA group had significantly higher seizure frequency (Welch’s t-test, n=8 per group, each point represents a mouse). **E**, Correlation between fostrocyte count and seizure frequency for each mouse. Statistical analysis compared fits to the data sets with either a horizontal line (null hypothesis) or a sloped line, finding that GNCTRL was significantly fit better with a sloped line (GNCTRL, N=6; GSDTA, N=7 mice, P=0.019).

Mice were then monitored for 6 weeks while seizures were recorded. Seizure frequencies in the GSDTA group were highly variable, so we binned the seizure results over a 3-day epoch (Fig. 5C). These results showed a large increase in seizure frequency in the GSDTA group that achieved statistical significance at later stages of the experiment. Comparison of the total seizure frequencies measured between days 6 and 40 showed a large increase in seizure frequency in the GSDTA group (Fig. 5D). Five mice injected with GSDTA had seizure frequencies greater than 5 Sz/D, which are much higher than we have ever observed in epileptic mice.^12^ Our validation studies show that seizures are required to trigger the appearance of TRAPed fostrocytes. Consistent with this observation, fostrocyte counts in the GNCTRL group showed a positive correlation with seizure frequency (Fig. 5E). In contrast, there was no correlation in the GSDTA group, where most of the hippocampal fostrocytes were ablated.

### 3.6 Testing the role of fostrocytes in chronic epilepsy using a paired experimental design

To increase rigor, we studied the role of fostrocytes using a cross-over design. We recorded baseline seizure frequencies for 2 weeks, injected 4OHT, and then recorded for 2 more weeks, which allows before-to-after paired comparisons (Fig. 6A). AAV mixtures were the same as above. All spontaneous seizures were similar in terms of their EEG (high amplitude, high frequency spiking, Fig. 6B, C), duration (40 s), and tonic-clonic motor components (Racine class 5; Supplemental Table 2). The GSDTA group showed a sharp increase in the number of seizures within days of 4OHT injection (Fig. 6D). Statistical analysis compared the seizure frequency of each mouse before and after 4OHT treatment. 4OHT had no effect in the GNCTRL group (Fig. 6E), but increased seizure frequency in the GSDTA group (Fig. 6F). Taken together, these results demonstrate that fostrocyte ablation increases seizure burden in epileptic TRAP2 mice.

**Fig. 6.**
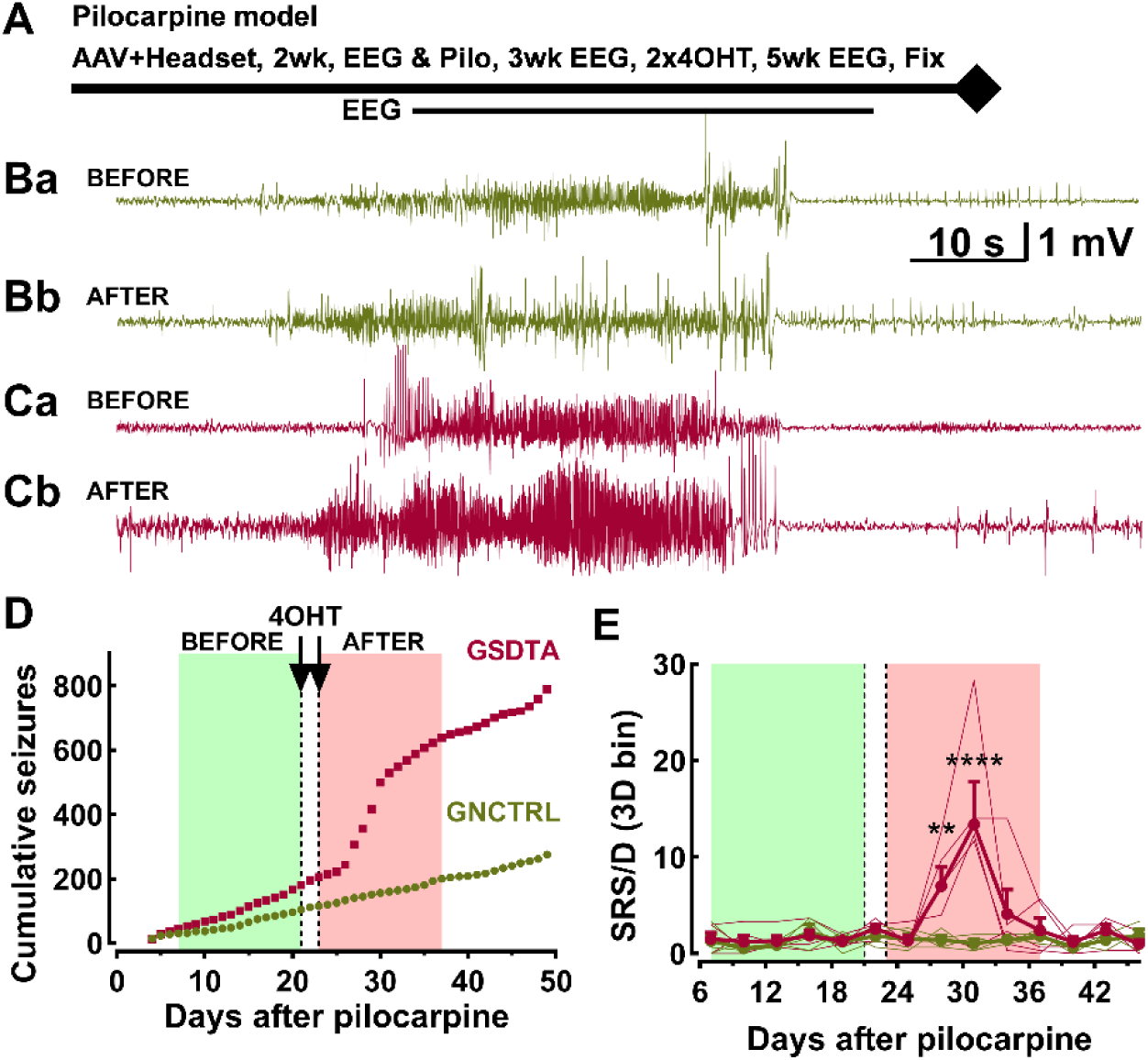
Testing the role of fostrocytes in chronic epilepsy using a paired experimental design. **A,** Schematic of experimental timeline. **B,** Representative traces EEG traces of convulsive seizures recorded before and after 4OHT treatment for the GNCTRL group before (Ba) and after (Bb) and the GSDTA group before (Ca) and after (Cb). **D,** Plot of the cumulative seizures for each group and the treatment periods. The EEG traces BEFORE 4OHT correspond to Day 19, while the traces AFTER 4OHT correspond to Day 34. **E**, Analysis of the average seizures per 3-day bin over the indicated 2-week windows. Statistical analysis by 2-way ANOVA and Fisher’s Least Significant Difference test. Asterisks represent the following P values: **, P<0.01; and ****, P<0.0001.

## 4. DISCUSSION

This study shows that seizures activate c-fos in astrocytes. Although c-fos expression in astrocytes was described in 1989,^33^ it is only recently that the importance of this signaling cascade in astrocytes has been established in disease,^22, 34^ ethanol consumption,^35^ and even learning.^23^ When seizures are induced in TRAP2 mice, c-fos expression is high enough to induce expression of Cre-ERT2, which in the presence of 4-hydroxy-tamoxifen, leads to excision of the floxed-stop cassette and expression of tdTomato.^4^ The TRAP2 system has been used successfully to map tdTomato+ neurons activated after SE and focal seizures.^19, 20^ While these studies noted the presence of tdT+ astrocytes, they were not quantified.^20^ Therefore, this is the first study to examine the role of c-fos activation in astrocytes after seizures. We established that these tdTomato+ glia were bona fide astrocytes using immunocytochemistry with the established astrocyte markers Sox9, S100B, and GFAP.^21^ The key conclusions from this study were that the vast majority of astrocytes transiently express c-fos after a discrete seizure, while a subset of astrocytes show prolonged expression of c-fos after SE. Importantly, in rodents where pilocarpine induces SE that lasts greater than 1 hour later develop spontaneous seizures that resemble human TLE seizures.^36^

To recapitulate, we define fostrocytes as a subset of astrocytes that are activated by seizures and that express c-fos to a level high enough to be labeled with tdTomato in TRAP2 mice. We show that fostrocytes are induced by seizures using two different chemoconvulsants, kainic acid and pilocarpine. We also show that fostrocytes are found in the hippocampus, following the same rank order of abundance in both models with CA1 ≥ CA3 > dentate gyrus. A limitation of these localization studies is that fostrocytes were often clumped together near the junction of the dentate and CA fields and often on the blood vessels that run in this hippocampal fissure, making it difficult to resolve individual cells. A similar clustering of c-fos+ astrocytes was also noted in a mouse model of experimental autoimmune encephalomyelitis.^34^ We emphasize that seizures are required to induce a state change in astrocytes to become fostrocytes. We used titrated doses of chemoconvulsants to induce SE, while avoiding mortality. As reported previously in rats,^37^ neuronal death occurred predominantly in the CA3 field, leading to hippocampal sclerosis and state changes in microglia and astrocytes, which obscured any direct effect of GSDTA on gliosis.

We show that fostrocytes display a reactive astrocyte morphology with shorter, thicker, and less complex processes. Reactive astrocytes can either be neurotoxic and contribute to the development of epilepsy,^38^ or neuroprotective.^39^ Our approach to infer the role of fostrocytes in epilepsy was to ablate them. To this end, we engineered novel Cre-dependent AAVs to express the A chain of diphtheria toxin (DTA). This chain acts enzymatically to ADP-ribosylate elongation factor 2, which stops protein translation, leading to apoptotic death.^27^ We re-engineered the AAV using a FLEX2 design with two key features: one, use of lox66 and lox71 mutated Cre binding sites^40^; and two, a split open-reading frame where only the amino-terminal fragment of DTA is in the DIO cassette (Supplemental Fig. 1). We show that co-injection of GSDTA and GFP AAVs successfully ablates this GFP signal. A possible limitation to the study is that DTA could affect neighboring cells. We measured the ability of GSDTA to ablate astrocytes in TRAP2 mice, finding a 26% decrease in Sox9+ labeled cells. Across the TLE models we find that fostrocytes make up 10-14% of hippocampal astrocytes. Taken together, we estimate off-target ablation of tdT-astrocytes at about 10%. We conclude that GSDTA specifically ablates fostrocytes in TRAP2 mice with minimal off-target effects on other astrocytes, neurons, or in the absence of Cre recombinase.

We investigated the role of fostrocytes on post-SE spontaneous seizures using two experimental designs: one, where we ablated fostrocytes 24 h after SE; and two, where we ablated fostrocytes after recording a baseline period. AAV injections and EEG headset surgeries were performed 2-3 weeks before pilocarpine-induced SE. We used both male and female mice in equal proportions that were randomized between groups, and all investigators were blind to the AAV injected. We used our optimized pilocarpine model where mice were treated with pilocarpine while on the EEG rig.^12^ The duration of SE was similar in both experiments (∼284 min), as was the time to the 1^st^ pilocarpine seizure and time to SE onset (Supplemental Table 1). This rules out the problem that SE duration affects seizure frequency.^36^ We also found that the time to the 1^st^ pilocarpine seizure and time to SE onset was not affected by the type of AAV injected. We highlight the AAV injection scheme as it differs from most AAV studies that only include GFP in their control group, including our early studies.^18^ The underlying assumption is that GFP is inert, when in fact it has toxic effects,^41^ which confounds any conclusions. In both experiments, mice injected with GSDTA and treated with 4OHT showed increased seizure frequencies. We conclude fostrocytes are playing beneficial roles in suppressing seizures, making them an attractive target for therapies.

Recent advances in astrocyte biology have led to paradigm shift in the roles they play, moving away from gluing neurons together to shaping behaviors such as memory. Simultaneously, the role of c-fos in astrocyte biology has expanded,^22^ as well as its expression in subclusters of astrocytes.^42^ Our analysis of the Seattle Alzheimer’s Disease Brain Cell Atlas revealed an upregulation of Fos and Jun RNA in multiple astrocyte clusters in AD patients with cognitive decline.^43^ RNA-seq studies from patients with Huntington disease and controls also found that clusters of astrocytes that expressed both *FOS* and *JUN* genes.^44^ CRISPRi screening of human astrocytes identified enhancers implicated in Alzheimer’s disease, leading to detailed regulatory map of the transcription factors activated by c-fos.^45^ Finally, our conclusion that fostrocytes play beneficial roles is supported by mouse studies showing that both *Fos* and *Jun* are expressed in anti-inflammatory astrocytes.^46^

## AUTHOR CONTRIBUTIONS

Madison Failor: Validation; supervision; formal analysis; writing—first draft; writing—review & editing. Bradley Bork: Validation; formal analysis. Jennifer San Pietro: Validation; formal analysis. Aleksandra Maciejczuk: Validation; formal analysis. Kendall Elmer: formal analysis. Corinne Lile-King: formal analysis. Mariia Oliinyk: formal analysis. Junguhk Choi: formal analysis. Andrew Huang: formal analysis; data curation. Dominik Tabor: formal analysis. Ronald Gaykema: Methodology; data curation; investigation; supervision. Edward Perez-Reyes: Conceptualization, supervision; methodology; formal analysis; writing—review & editing.

## ACKNOWLEDGEMENTS

We thank Dr. Jaideep Kapur for helpful discussions and for donating homozygous TRAP2 mice. We thank Ashley Brandebura and Leon Straub for comments on the manuscript. This research was funded by a Harrison Undergraduate Research Award (B.B.), a Double Hoo Award (M.J.F. & J.M.S.P.), and National Institutes of Health grants NS112549 (E.P.R.), MH135905 (E.P.R.), U19AG060909 (SEA-AD), and T32 GM148379 (M.J.F.).

## CONFLICT OF INTEREST STATEMENT

The authors declare no conflict of interest.

## ETHICS STATEMENT

We confirm that we have read the Journal’s position on issues involved in ethical publication and affirm that this report is consistent with those guidelines.

## DATA AVAILABILITY STATEMENT

The data presented in this study are available in this article and upon request.

## Notes

### Competing Interest Statement

The authors have declared no competing interest.

