## Supplemental Fig. 1 for "Seizures induce c-fos expression in a subset of astrocytes, termed fostrocytes, that dampen subsequent seizures"

**A**

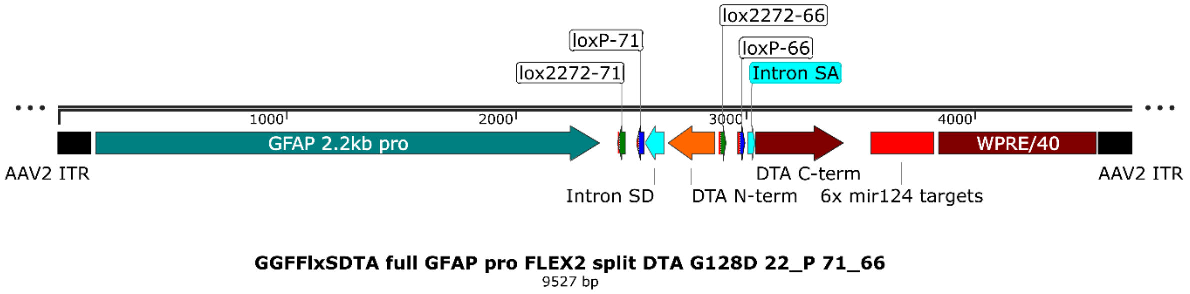


**B**

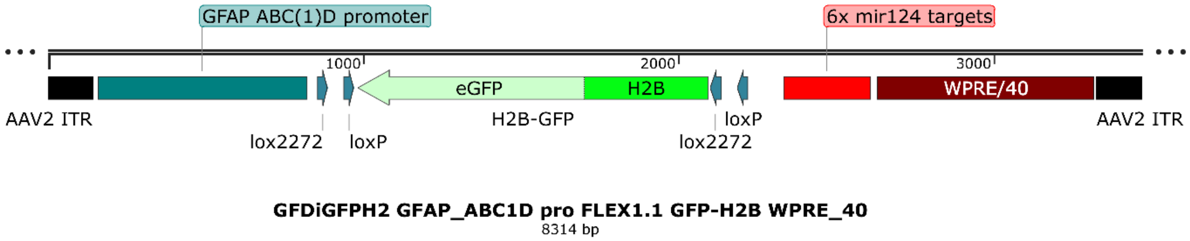


**C**

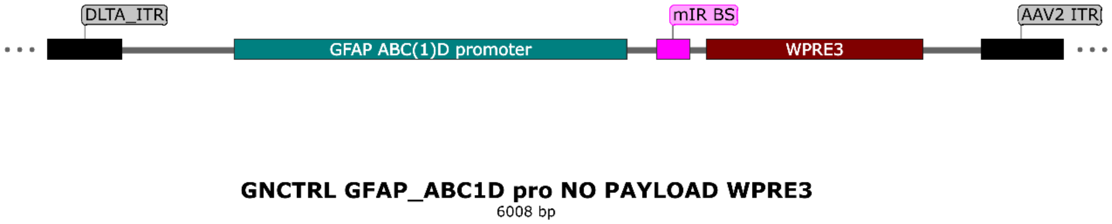


**D**

| AAVs | Stock (E13) | Titer inj (E13) | MZ | MN | VG injected (E10) |
|---|---|---|---|---|---|
| AAV5/GLFFlxSDTA | 5.17 | 0.50 | 14.3 | | 1.5 |
| AAV5/GFDiGFPH2 | 22.30 | 0.50 | 3.3 | | 1.5 |
| PBS (+Ca/Mg) | | | 130.0 | | |
| AAV9/GNCTRL | 3.30 | 0.50 | | 22.5 | 1.5 |
| AAV5/GFDiGFPH2 | 22.30 | 0.50 | | 3.3 | 1.5 |
| PBS (+Ca/Mg) | | | | 122.0 | |

**Supplemental Fig. 1 Description of AAV constructs engineered to study fostrocytes**. **A**, a Cre-dependent AAV construct that delivers a split diphtheria toxin A chain (DTA). This construct uses the 2.2-kb GFAP promoter (a gift from Karl Deisseroth, Addgene plasmid 27055). The Cre recombinase binding sites of the DIO/FLEX switch carry the low recombination mutations 66 and 71 (Albert et al., 1995; Fischer et al., 2019). The DTA harbors the G128D attenuation mutation (Maxwell et al., 1987). The DTA coding region was split at R77 to disrupt the ADP-ribosylation catalytic site (Rodnin et al., 2020). The AAV also contains 6 copies of mir124 target sequences to reduce off-target expression in neurons (Taschenberger et al., 2017). The 3'-UTR is a truncated WPRE sequence combined with a minimal SV40 pA signal (Choi et al., 2014). **B**, a Cre-dependent AAV construct that delivers an H2B-GFP fusion protein. This construct uses the 681-bp truncated GFAP promoter GfaABC$_1$D (Lee et al., 2008). The DIO/FLEX switch uses the original loxP/lox2272 spacers with wild-type Cre binding sites (Schnutgen and Ghyselinck, 2007). **C**, Empty payload AAV used as negative control that contains a similar promoter and 3'-UTR but does not lead to the expression of any gene product. **D**, Key used for blinding the AAV injected. Plan includes the titer of both stocks and dilutions, and the number of AAV vector genomes (VG) injected per mouse. Viral aliquots were labeled MZ and MN, where MZ contains SDTA and GFP, while MN contains GNCTRL and GFP. Note that each injection delivers the same amount of GFP and total AAV vector genomes.
